# Shallow-angle intracranial cannula for repeated infusion and in vivo imaging with multiphoton microscopy

**DOI:** 10.1101/2025.01.22.634409

**Authors:** Steven S. Hou, Joyce Yang, Yeseo Kwon, Qi Pian, Yijing Tang, Christine A. Dauphinais, Maria Calvo-Rodriguez, Mirna El Khatib, Sergei A. Vinogradov, Sava Sakadzic, Brian J. Bacskai

## Abstract

Multiphoton microscopy serves as an essential tool for high-resolution imaging of the living mouse brain. To facilitate optical access to the brain during imaging, the cranial window surgery is commonly used. However, this procedure restricts physical access above the imaging area and hinders the direct delivery of imaging agents and drugs. To overcome this limitation, we have developed a cannula delivery system that enables the implantation of a low-profile cannula nearly parallel to the brain surface at angles as shallow as 8 degrees, while maintaining compatibility with multiphoton microscopy. To validate this approach, we perform direct infusion and imaging of various fluorescent cell markers in the brain. Additionally, we successfully demonstrate tracking of degenerating neurons over time in Alzheimer’s disease mice using Fluoro-Jade C. Furthermore, we show longitudinal imaging of brain tissue partial pressure of oxygen using a phosphorescent oxygen sensor. Our developed technique should enable a wide range of new longitudinal imaging studies in the mouse brain.

## INTRODUCTION

*In vivo* imaging with multiphoton microscopy has become an indispensable technique for studying cortical function in the intact brain (Helmchen & Denk, 2005; Lecoq et al., 2019; Trachtenberg et al., 2002). Due to the opacity of the mouse skull, a crucial step in preparation for multiphoton imaging is the surgical procedure that allows optical access to the brain. In the thinned skull approach, a small portion of the skull is made transparent by carefully drilling down to ∼20 µm (Drew et al., 2010; G. Yang et al., 2010). While this approach is minimally invasive, the skull regrowth limits its longitudinal application. In the cranial window approach, a bone flap from the skull is completely removed and a cover glass is sealed over the exposed brain (Cramer et al., 2021; Holtmaat et al., 2009). While the cranial window grants optical access to the brain, physical access to the brain is restricted.

The delivery of chemical compounds, including imaging agents and drugs, into brain is an essential component of many multiphoton microscopy imaging studies. However, many compounds cannot cross the blood-brain-barrier (BBB) and therefore, cannot be delivered to the brain through systemic administration (Banks, 2009). To enter the brain, they need to be directly applied, either topically to the brain surface or by injection with a needle or micropipette (Hillman, 2007). Since the cranial window imposes a physical barrier to the brain, compounds often need to be delivered to the brain at same time as cranial window surgery, and before the cover glass is sealed. Since further access to the brain is restricted after the cranial window is in place, the ability to repeatedly deliver to brain and perform longitudinal imaging is not possible using the standard cranial window approach.

Recently, new techniques have been developed that combine direct delivery to the mouse brain with *in vivo* imaging capability. One class of methods uses soft, transparent materials in place of the cover glass for the cranial window (Heo et al., 2016; N. Yang et al., 2022). The softness of the material permits penetration of needles and micropipettes, while its transparency is sufficient for multiphoton microscopy. A related technique incorporates silicone access ports within the glass cover slip, enabling injections through these ports while simultaneously allowing for imaging through the cover glass (Roome & Kuhn, 2014). Another approach to provide access to the brain has been the development of removable cranial windows (Goldey et al., 2014). Here, the cover glass can be removed to grant temporary access to the brain and then resealed. The main drawback of these techniques is that they require the repeated insertion of the injection needle or micropipette into the brain tissue which may lead to tissue damage, bleeding and inflammation. This can significantly affect the quality of the imaging and impede longitudinal imaging. Furthermore, the removal of the cranial window can cause additional inflammation from recurring brain exposure.

Techniques based on implantable devices have also been used for delivery into brain. In one approach, a micro-optical fluidic device is created with an inlet and outlet for infusion and removal of chemical compounds (Takehara et al., 2014). While this approach allows repeated application of compounds at the brain surface, the relative amounts of the compound that reaches different depths of the brain parenchyma cannot be varied and are strongly dependent on its diffusion properties. Also, the need for dura removal during implantation of the device could lead to complications after surgery recovery. In another technique, commercially available cannulas are implanted into the brain for delivery while cranial windows are placed adjacent to the cannula (Zhang et al., 2019; Zuluaga-Ramirez et al., 2015). However, the larger size of these cannulas imposes constraints on the implantation to steeper angles (previous studies have used 45 degrees (Zhang et al., 2019) and 90 degrees (Zuluaga-Ramirez et al., 2015)). For cannulas targeted towards the superficial cortex, the steeper angles result in the cannula outlet location being close to the edge of the cranial window, where imaging can be challenging and only a small part of the imageable brain area may be exposed to the injected substance.

In this study, we developed a custom cannula delivery system that enables the permanent implantation of a low-profile cannula at angles as shallow as 8 degrees into the mouse brain. The shallow angle of the implanted cannula allows its tip to be centered over a 4 mm cranial window while still being located within the superficial layers of the mouse cortex. Importantly, the cannula does not interfere with the cranial window or with imaging using multiphoton microscopy. We validated our method using various fluorescent cell markers to confirm successful delivery and imaging capability. Additionally, we demonstrated repeated infusion and longitudinal imaging by tracking neurodegeneration in a mouse model of Alzheimer’s disease using Fluoro Jade C, a fluorophore traditionally employed for staining *ex vivo* tissue sections (Schmued et al., 2005). Furthermore, we demonstrated longitudinal functional imaging of tissue partial pressure of oxygen (pO_2_) through direct infusion of the phosphorescent oxygen sensor, Oxyphor 2P (Esipova et al., 2019) and *in vivo* phosphorescence lifetime imaging (Finikova et al., 2008; Parpaleix et al., 2013; Sakadžić et al., 2010).

## RESULTS

### Development of custom cannula, holder and procedure for shallow-angle implantation

Our objective was to develop a cannula-based delivery method for repeated infusion into the mouse cortex while maintaining the ability to perform *in vivo* multiphoton microscopy through a cranial window. This combined cannula and cranial window system needed to meet several criteria: 1) The cranial window needed to be of adequate size (4 mm) to image a relatively large portion of the underlying cortex. 2) The cannula’s tip had to be centered relative to the cranial window. 3) The cannula’s outlet had to be within the superficial to deep layers of the cortex, which are regions that are directly imageable by multiphoton microscopy. These requirements necessitated designing a low-profile cannula that could be implanted at shallow angles relative to the brain surface, ensuring the cannula would not obstruct the placement of the cranial window cover glass (**Fig. 1a, b**).

**Figure 1.**
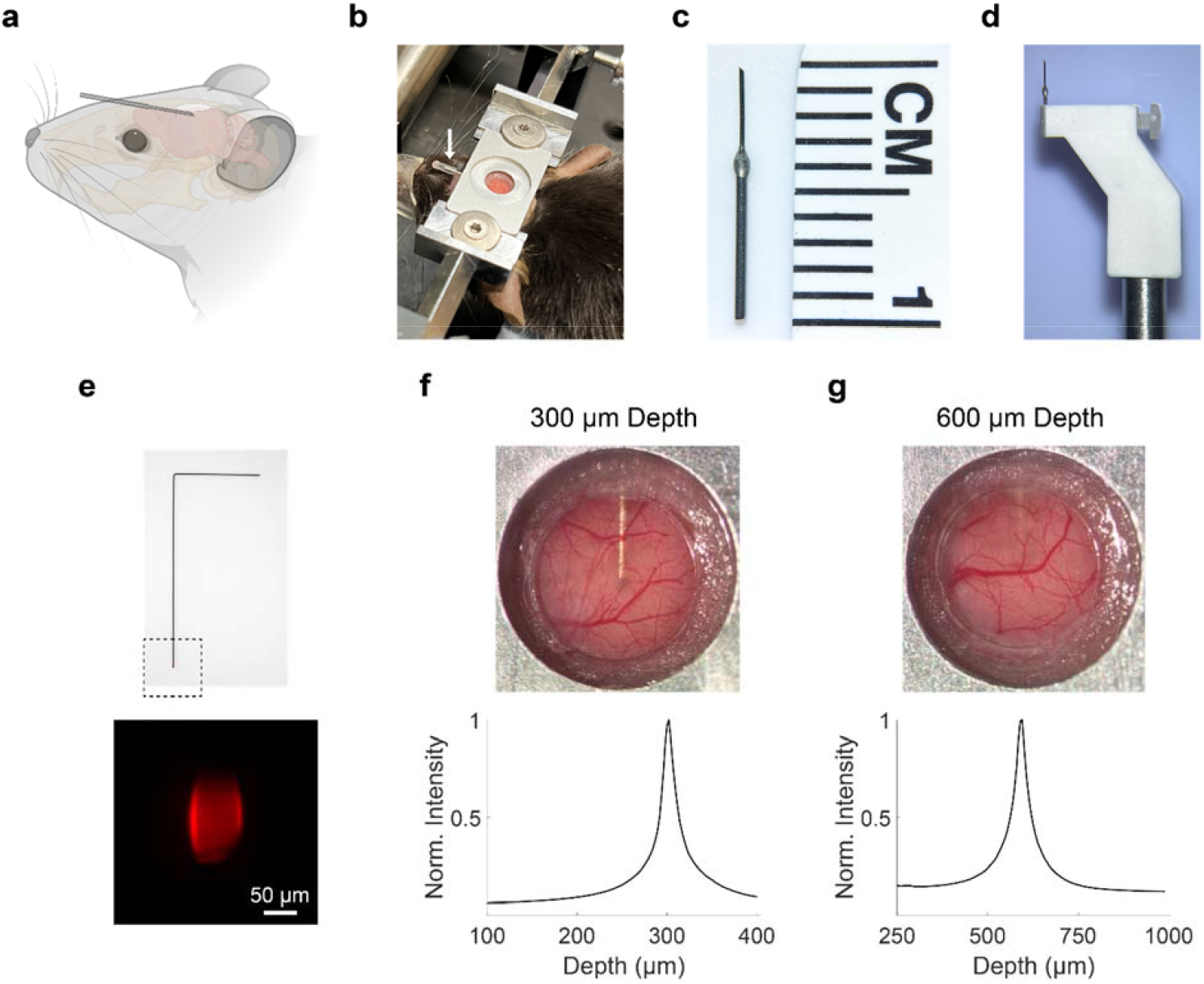
Design and characterization of shallow-angle cannula delivery system. **(a)** Schematic showing the position and the shallow approach angle of the implanted cannula in the mouse brain. (**b**) Image showing implanted cannula in a C57BL/6J mouse. White arrow indicates location of the implanted cannula. The cannula tip is sealed by an attached segment of polyethylene tubing with a mixture of dental acrylic and cyanoacrylate glue at its end. (**c**) The cannula is created by combining segments of 26-gauge and 33-gauge stainless steel tubing. (**d**) A custom cannula holder secures the cannula during implantation. (**e**) A fluorescent dummy wire is created to determine the depth of the implanted cannula. The length of the dummy wire is matched to the length of the cannula with the tip of the wire coated with SR101. Images of the cranial window after cannula implantation at 300 µm (**f**) and 600 µm (**g**) depths. The cannula is clearly visible at 300 µm depth and a faint shadow of the cannula is visible at 600 µm depth. To confirm the location of the cannula tip, the fluorescent dummy wire was inserted into the cannula and multiphoton microscopy was used to determine the depth-dependent fluorescence intensity profile.

The cannula was created by combining segments of 26-gauge and 33-gauge stainless steel tubing (**Fig. 1c**). The portion of the cannula to be implanted inside the brain tissue was chosen to be 33-gauge to minimize impact on the brain tissue. A 45-degree bevel was added to the cannula tip to facilitate puncturing of the dura during implantation. Due to the fragility of the 33-gauge tubing, the portion of the cannula outside the brain was reinforced by the larger diameter 26-gauge tubing. A custom holder was created to securely hold the 26-gauge portion of the cannula (**Fig. 1d**). The dimensions of the holder were chosen so that the holder would not make contact with the mouse head during the implantation process.

We tested two depths for cannula implantation: 300 µm and 600 µm. At 300 µm depth, corresponding to an implantation angle of 8 degrees, the entire outline of the cannula up to its tip was visible in the cranial window image (**Fig. 1f**). At 600 µm depth, corresponding to an implantation angle of 16 degrees, only a shadow of the cannula was visible at the top of the cranial window (**Fig. 1g**). We devised a method to verify the depth of the cannula after implantation by creating fluorescent dummy wires made of tungsten (**Fig. 1e**). The dummy wires, matched in length to the cannula, had tips coated with the red fluorophore, sulforhodamine 101 (SR101). After inserting the dummy wire into the implanted cannula, we imaged the fluorescence in a volume surrounding the cannula tip. By plotting the fluorescence intensity as a function of depth, we could confirm the accurate localization of the cannula tip for both 300 µm and 600 µm depths (**Fig. 1f, g**).

We evaluated the neuroinflammatory response to the chronically implanted cannula in the brain by immunostaining for microglia and activated astrocytes in both the ipsilateral cortical region (right hemisphere, site of cannula implantation and cranial window) and the contralateral cortical region (left hemisphere, control). To better understand the direct effect of cannula implantation, we also evaluated neuroinflammation in mice with cranial windows alone. At 30 days post-surgery, cannula-implanted mice showed a localized increase in microglia and astrocyte activation directly proximate to the location of the cannula and sparse astrocyte activation in other areas of the ipsilateral hemisphere, away from the cannula site (**Fig. S3b**). In cranial window mice, we also observed sparse activation of astrocytes in the ipsilateral hemisphere (**Fig. S3a**). We did not observe activated astrocytes in the contralateral hemisphere for either the cannula or cranial window mice. We performed quantitative analysis of microglia and activated astrocytes density in regions excluding the cannula site. For microglia density, we found that there were no significant differences between the ipsilateral and contralateral hemispheres in both the cranial window and cannula-implanted mice (**Fig. S3c**). Furthermore, a comparison of activated astrocyte density in the ipsilateral hemisphere between cranial window alone and cannula mice also showed no significant differences (**Fig. S3d**).

### Infusion and imaging of various cell type specific markers, structural markers and markers for Alzheimer’s disease pathology in the mouse brain

We validated the ability of our approach to label and image various cellular targets *in vivo* by infusing a wide assortment of fluorescent cell markers and imaging with multiphoton microscopy. Our tests included both general cellular imaging markers and markers for imaging pathological processes in the brain. We specifically focused on fluorescent markers known to be challenging to introduce into the brain after systemic delivery due to their inability to cross the BBB. Many of the fluorophores we tested have been extensively used for *in vitro* live cell imaging experiments but previously have limited application for *in vivo* brain imaging.

We infused a nuclei acid binding fluorophore, Hoechst 33342 and observed bright and dense labeling of nuclei in the brain region surrounding the cannula outlet location (**Fig. 2a**). Following infusion of NeuroTrace 500/525 (Damisah et al., 2017) and intravenous (IV) injection of Fluorescein Dextran, we could detect labeled pericytes in close proximity to blood vessels. (**Fig. 2b**). We could successfully image astrocytes (**Fig. 2c**) and microglia (**Fig. 2d**) after infusion of SR101 (Nimmerjahn et al., 2004) and Isolectin GS-IB4, Alexa Fluor 594 Conjugate (Eichhoff et al., 2011) respectively. We could visualize and track individual lysosomes after infusion of LysoTracker Green DND-26 into the brain (**Fig. 2e**). By infusing Fluorescein labeled Wisteria Floribunda Lectin (WFL), we specifically targeted perineuronal nets (PNNs) which are extracellular structures surrounding inhibitory neurons (Brückner et al., 1996). Using this approach, we found dense labeling of PNNs throughout the cannula outlet area (**Fig. 2f**).

**Figure 2.**
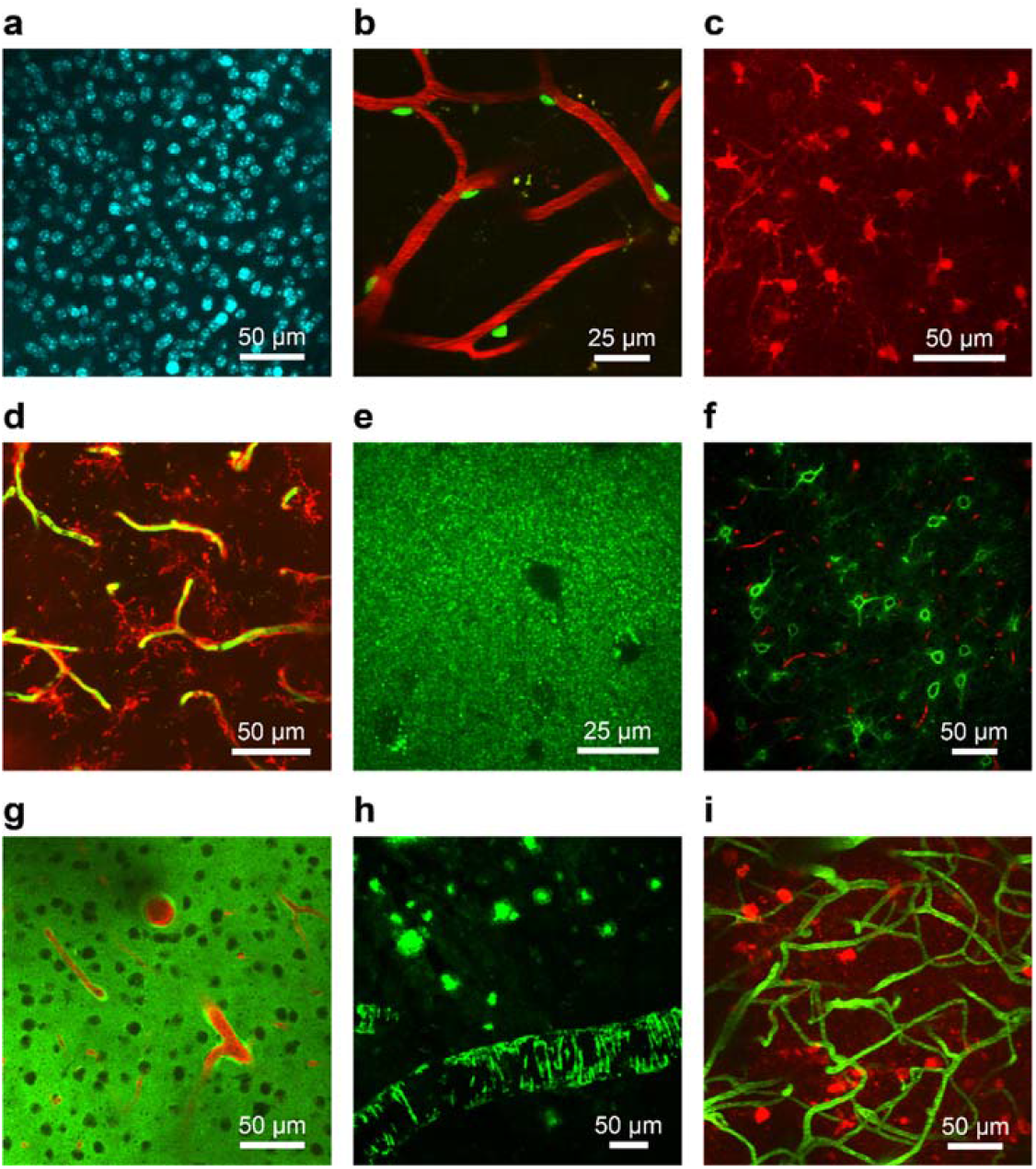
Validation of cannula delivery system using various fluorescent cell markers and probes for AD pathology. Fluorophores were infused through the cannula system and multiphoton microscopy was used for *in vivo* imaging. Images are shown as maximum intensity projections of 3D volumes. Each fluorescent cell marker was delivered to n = 3 C57BL/6J mice. **(a)** Cell nuclei were labeled after infusion of Hoechst 33342. (**b**) Pericytes (green) were labeled after infusion of NeuroTrace 500/525 and Rhodamine B Dextran (red) was injected IV to act as an angiogram. (**c**) Astrocytes were labeled following the infusion of SR101. (**d**) Microglia (red) were labeled after infusion of Isolectin GS-IB4, Alexa Fluor 594 Conjugate. The vasculature was labeled using IV injection of Fluorescein Dextran (green). (**e**) Lysosomes were labeled after infusion of LysoTracker Green DND-26. (**f**) Perineuronal nets (green) were labeled after infusion of Fluorescein labeled WFL and Rhodamine B Dextran (red) was injected IV. (**g**) Fluorescent Aβ42 (green) was infused to demonstrate perivascular clearance out of the brain (n = 3 C57BL/6J mice). Rhodamine B Dextran was injected to label the vasculature (red). (**h**) ThioS was infused and showed labeling of amyloid plaques and CAA in APP/PS1 mice (n = 3). (**i**) Thiazine Red was infused in rTg4510 mice (n = 3) and showed labeling of neurofibrillary tangles (red). Fluorescein Dextran (green) was injected IV to serve as an angiogram.

To demonstrate the utility of our approach for studying disease models in the brain, we also tested the infusion of fluorophores that could be applied for Alzheimer’s disease (AD) imaging. We explored the distribution of the amyloid-β (Aβ) peptide in the brain by direct infusion of fluorescently labeled Aβ42 peptide (HiLyte Fluor 488, Aβ42) into the mouse cortex. Rhodamine B Dextran was systemically injected to create an angiogram. In regions above the infusion location, we observed accumulation of fluorescent Aβ around arterioles and arteries (**Fig. 2g**). This perivascular fluorescence is consistent with previous reports after acute intracranial injection of fluorescent tracers (Iliff et al., 2012). Additionally, we tested our technique for imaging AD pathology in two mouse models of AD. As a model of cerebral amyloidosis, the APP/PS1 mouse model was used, which develops amyloid pathology starting at 5-6 months of age (Jankowsky et al., 2004). For labeling of pathology, we tested Thioflavin S (ThioS), a fluorescent dye considered the gold standard for *ex vivo* labeling of amyloid aggregates (Rajamohamedsait & Sigurdsson, 2012). After infusing ThioS in APP/PS1 mice, we observed strong *in vivo* labeling of amyloid plaques and cerebral amyloid angiopathy (CAA) in the brain (**Fig. 2h**). We used the rTg4510 mouse model as a model of tauopathy, which starts development of neurofibrillary tangles (NFTs) at 2 months of age (Santacruz et al., 2005).

Following infusion of Thiazine Red, a fluorescent dye known to bind to NFTs *ex vivo* (Rudinskiy et al., 2014), we found that we could successfully label and image NFTs in the brains of rTg4510 mice (**Fig. 2i**).

### Longitudinal tracking of neurodegeneration in a mouse model of Alzheimer’s disease

We next used our cannula-based delivery method to perform longitudinal imaging in the mouse brain. Our aim was to develop a technique to track neurodegeneration in real time in the mouse brain. To identify degenerating neurons, we used the fluorescent dye, Fluoro Jade C (FJC) which has traditionally been applied for the staining of tissue sections. We used rTg4510 mice to model neurodegeneration, as neuron loss in the cortex of these mice has been well-documented (Santacruz et al., 2005). We infused FJC weekly for 3 weeks and conducted imaging with multiphoton microscopy. Prior to each imaging session, we injected IV Rhodamine B Dextran to form a fluorescent angiogram. We used the vascular structure from the angiogram as a reference to acquire the same fields of view on a weekly basis. After infusion of FJC into the transgenic mice, we found cells labeled by FJC scattered throughout the brain (**Fig. 3**).

**Figure 3.**
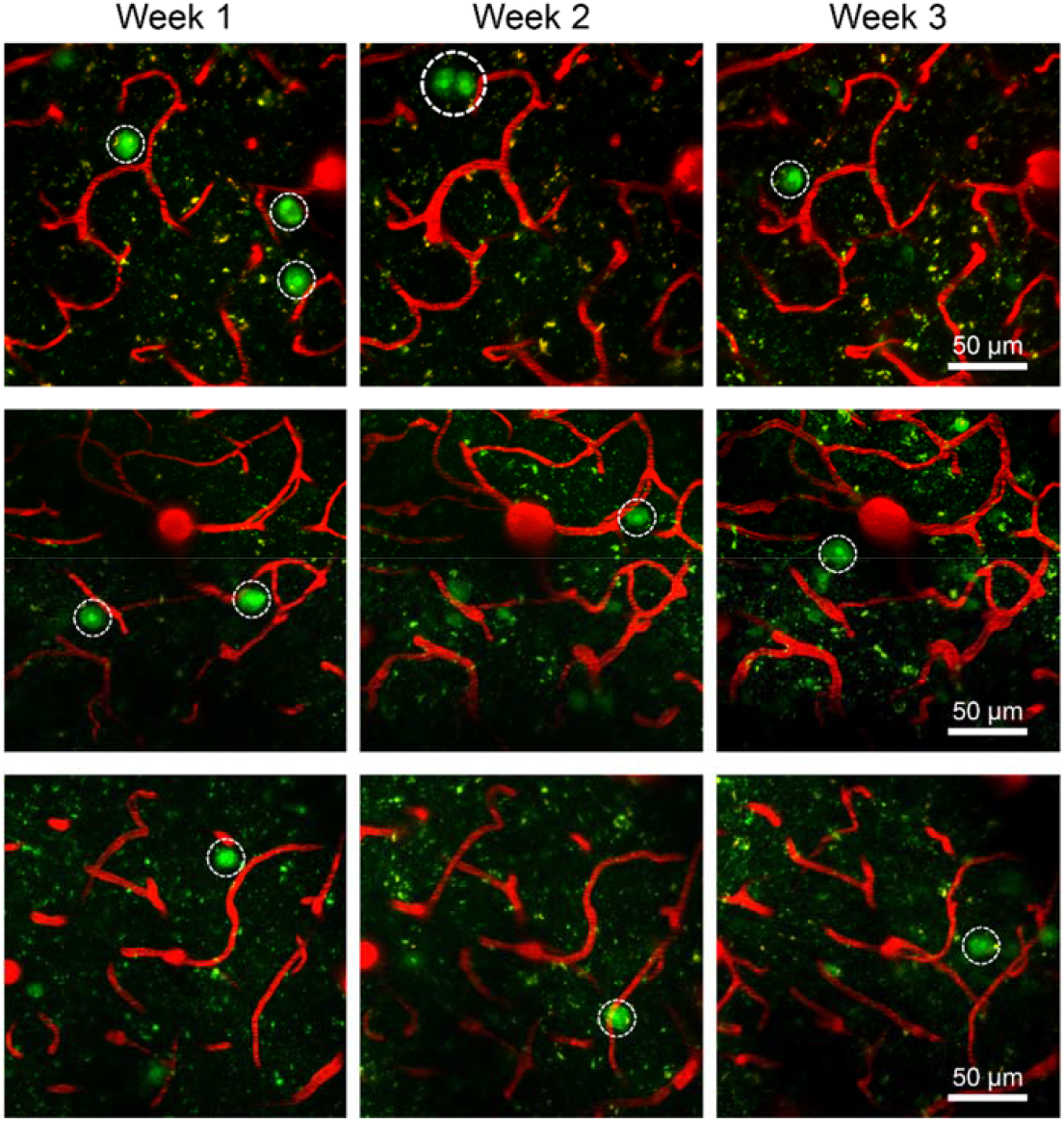
Longitudinal imaging of degenerating cells using Fluoro Jade C in rTg4510 mice. Fluoro Jade C (FJC, green) was infused through the cannula system on a weekly basis in rTg4510 mice (n = 3). The cannula was implanted at 300 µm depth. Rhodamine B Dextran (red) was systemically injected to create a fluorescent angiogram. Representative images show the appearance and disappearance of FJC positive cells during weekly imaging (FJC positive cells circled in white). Each row shows images from a unique mouse.

When comparing labeled cells in the same field of view, we found a dynamic process where cells labeled one week were mostly unlabeled the following week. Additionally, regions without labeled cells in one week sometimes displayed new labeled cells in the subsequent week.

When FJC was infused in age-matched control mice, we observed no FJC-labeled cells in the brain (**Fig. S1**). Our results indicate that our approach can successfully track neurodegeneration in a longitudinal manner in transgenic mice.

### Longitudinal imaging of tissue pO_**2**_ **in the awake mouse cortex**

We next applied our technique for longitudinal imaging to the measurement of tissue partial pressure of oxygen (pO_2_) in the brain. For measuring pO_2_ *in vivo*, we used the two-photon excitable phosphorescent oxygen sensor Oxyphor 2P (Esipova et al., 2019). The measured phosphorescence lifetime of the sensor can be directly mapped to the oxygen concentration in the brain. We infused Oxyphor 2P through our cannula delivery system on a weekly basis and collected two-photon phosphorescence emission data using a custom microscope. Fluorescein Dextran was injected IV to act as an angiogram. Weekly imaging was performed in awake mice. The imaging field was focused around diving arteries in the brain, where we expected to observe the highest contrast in pO_2_ values. Phosphorescence lifetimes were measured within a ∼210 µm region of the ∼350 µm imaging field at points in a 20×20 grid.

After infusion of Oxyphor 2P, we obtained high-contrast phosphorescence images of the probe down to 400 µm from the surface of the brain (**Fig. 4a**). Color-coded pO_2_ levels (in mmHg) were overlaid onto the phosphorescence intensity image at each grid point. Cell bodies were clearly visible in the intensity images due to the negative contrast they provided in the images. The consistently sharp images of the cells over time indicated that the spatial resolution of the imaging was not compromised by repeated infusion. As expected, we found elevated pO_2_ in regions close to the diving arteries and its branches and more uniform reduced pO_2_ values in surrounding regions. We also found that the measured pO_2_ spatial maps were consistent on a week-to-week basis, indicating that our infusion technique is viable for longitudinal tracking of pO_2_. Furthermore, we demonstrated that our technique allows imaging of pO_2_ in anesthetized mice (**Fig. S2**). In anesthetized mice, we generally observed higher pO_2_ values both close to diving arteries and surrounding regions compared to awake mice. We attribute this observed difference in pO_2_ levels to a reduction in local oxygen consumption in anesthetized mice. To determine the dependence of pO_2_ on depth in awake mice, we obtained the histogram for each depth and found that there is a shift of the pO_2_ distribution towards higher pO_2_ values at larger depths (**Fig. 4b**). Additionally, we measured the average of the bottom 20% of the pO_2_ distribution, corresponding to regions of the field of view away from the diving artery (**Fig. 4c**). We found an increase in the bottom 20% of the pO_2_ distribution for larger depths, which is consistent with a recent study measuring the depth dependence of pO_2_ in awake mice at a single time point (Mächler et al., 2022).

**Figure 4.**
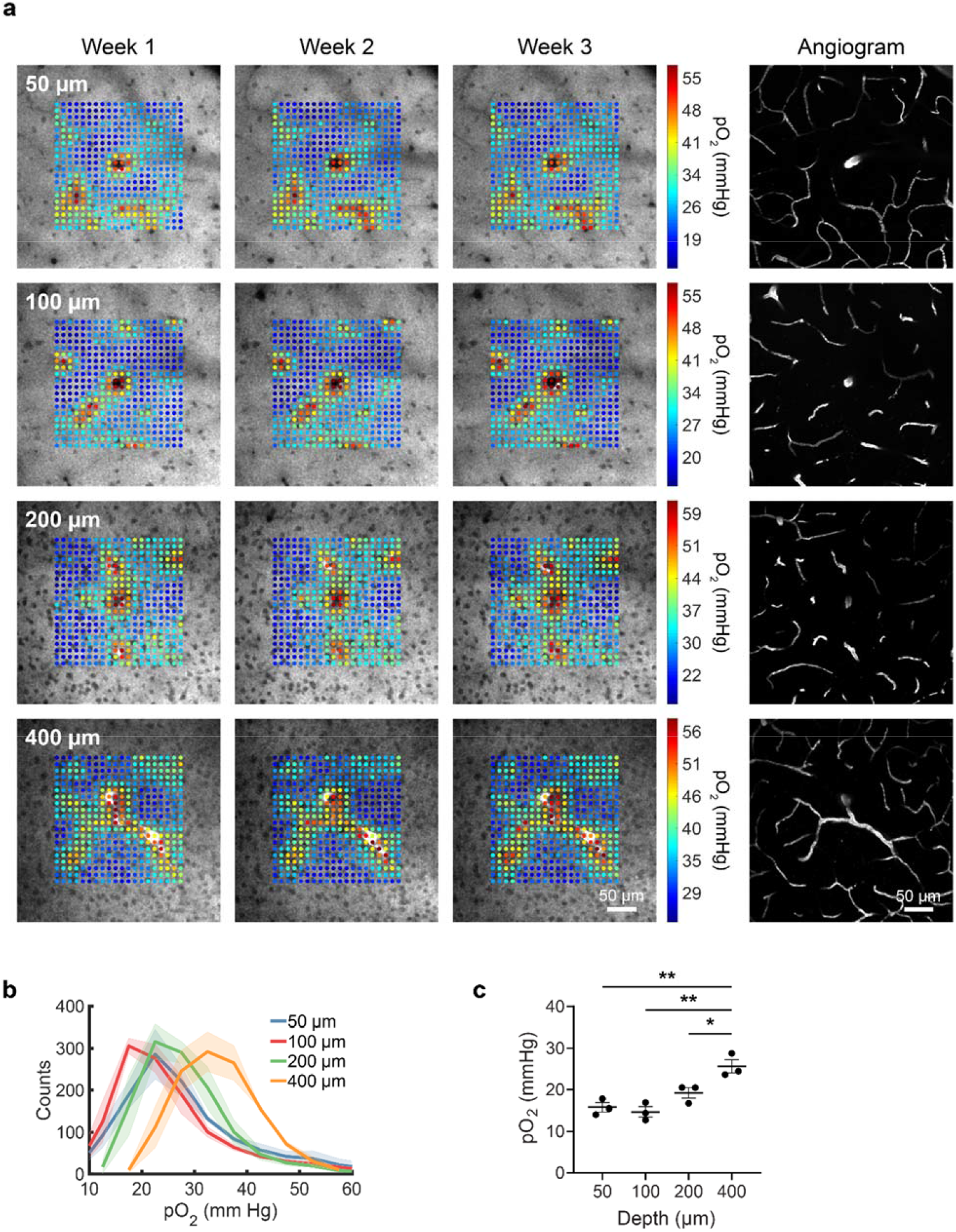
Longitudinal imaging of brain tissue pO_2_ in awake mice. Oxyphor 2P was infused through the cannula system on a weekly basis in C57BL/6J mice (n = 3). The cannula was implanted at 600 µm depth. Imaging was performed using multiphoton phosphorescence lifetime microscopy. During the first weekly imaging session, the field of view was selected around diving arteries. **(a)** Representative images show color coded pO_2_ values overlaid on top of phosphorescence intensity images. Images were acquired at 50, 100, 200 and 400 µm from the brain surface. Angiograms were acquired after IV injection of Fluorescein Dextran. The consistency and high contrast of the images from week to week demonstrate the viability of our approach for longitudinal pO_2_ imaging. (**b**) Histograms of pO_2_ values at each depth from n = 3 C57BL/6J mice. The pO_2_ values are based on the average of measurements from all three weeks. (**c**) Quantification of the bottom 20% of the pO_2_ distribution (averaged over three weeks) at each depth from n = 3 mice. Significance was calculated using one-way ANOVA with Tukey’s multiple comparison test: \**P* <0.05, \*\**P* <0.01.

## DISCUSSION

We have developed a cannula-based system that enables intracranial delivery into mouse brain and is fully compatible with *in vivo* multiphoton microscopy. The basis of our technique is that to achieve targeting of the superficial mouse cortex (∼300 µm) while keeping the cannula outlet centered relative to the cranial window necessitates a very shallow angle of approach. This led to the development of a low-profile cannula and an implantation procedure that is compatible with the shape of the mouse head. Importantly, since the angle of the cannula relative to the brain surface is nearly parallel, the cannula does not obstruct the imaging objective for multiphoton microscopy. This is in contrast to other cannula-based approaches for brain delivery which use bulkier commercial cannulas that occupy more space above the brain and are limited in how shallow the implantation angle can be.

Since the cannula in our approach is permanently implanted, it can bypass some of the issues associated with using soft, penetrable materials either as ports for the injection or for the entire cranial window. In these approaches, an injection needle or micropipette is inserted each time for brain delivery. This has potential to puncture blood vessels and create new tracts of tissue damage during each injection. This issue is especially significant when targeting of deeper regions of the brain. Additionally, the location for each injection may differ, depending on the injection accuracy. In our cannula-based technique, the damage to brain occurs once during the initial implantation. Also, infusion using the cannula is highly reproducible, as the outlet location of the cannula is stable. This ensures that the infused substance exposes the same brain regions at the same dosage, which makes comparisons after each infusion more straightforward.

Our technique may also offer certain advantages over traditional acute intracranial injections. In our approach, the cannula outlet is below the area of imaging while for acute injections the damage occurs from the brain surface down to the depth of the injection with the damaged region located centrally relative to the imaging area. Another key difference is that with our approach the infusion is performed when the brain has already recovered from the cannula implantation. In contrast for acute injections, the injection is performed immediately after the needle is inserted into the brain. This difference may influence the brain’s inflammatory state and its reaction to the delivered substance, particularly in the context of delivering viral vectors.

When we conducted pilot tests on our cannula-based system by infusing various imaging agents, we found that the infusion would sometimes cause neuroinflammation and an infiltration of immune cells into the brain, especially near the top of the brain. This would lead to the cranial window becoming more opaque and would ultimately limit the achievable depth of imaging. We found that we could frequently mitigate this issue by reducing the concentration of the imaging agent. Future studies need to be conducted to understand why certain chemical compounds trigger a response while others do not and how this response is influenced by infusion parameters including volume, concentration and rate.

We believe our developed technique will enable many new types of longitudinal imaging studies in the brain. So far, we have successfully demonstrated the tracking of neurodegeneration and the measurement of brain tissue partial pressure of oxygen over time. Given the wide assortment of imaging agents that are commonly used *in vitro* for live cell imaging, which have previously never been applied for *in vivo* imaging with multiphoton microscopy, we believe our approach should allow the testing of many of these agents *in vivo* and enable their application for longitudinal imaging. Additionally, our technique should have utility in preclinical imaging studies for the repeated delivery and imaging of therapeutic agents in the brain.

## METHODS

### Animals

We used C57BL/6J mice (The Jackson Laboratory) for validating our cannula-based delivery system using various fluorescent cell markers and for *in vivo* longitudinal pO_2_ imaging. Mice of both sexes at 4-6 months of age were used. For imaging of neurofibrillary tangles and tracking of neurodegeneration, we used rTg4510 (The Jackson Laboratory, Tg(Camk2a-tTA)1Mmay Fgf14Tg (tetO-MAPT*P301L)4510Kh) mice expressing the tetracycline-controlled transactivator protein (tTA) (Santacruz et al., 2005) along with age-matched non-transgenic littermates. Mice of both sexes at 6–8 months of age were used. For imaging of amyloid aggregates, we used APP/PS1 (The Jackson Laboratory, B6.Cg-Tg (APPswe, PSEN1-dE9)85Dbo/Mmjax) double transgenic mice expressing mutant human APPswe/human PS1-ΔE9 (Jankowsky et al., 2004). Mice of both sexes were used at 12-13 months of age.

Animals were housed in an AAALAC-certified facility (Massachusetts General Hospital, D16-00361), and all animal studies were performed under the supervision of the Massachusetts General Hospital Institutional Animal Care and Use Committee in accordance with the approved institutional protocol (2018N000131).

### Cannula, dummy wire and holder design

The custom cannula was constructed by combining a 6 mm length, 26-gauge stainless steel tubing (6689, New England Small Tube) with a 9.6 mm length, 33-gauge stainless steel tubing (33RW-378-45deg, Vita Needle). The tip of the 33-gauge tubing was beveled at 45-degree. The 26-gauge tubing was aligned to the 33-gauge tubing using a custom cannula assembly part and the two tubings were attached using epoxy (ClearWeld, J-B Weld). For cannulas used in this study, the cutting, beveling and assembly was also performed at Protech International. The cannula holder was machined from high-density polyethylene and a hole was created using a 0.0189” drill bit (McMaster-Carr) for securing the 26-gauge tubing (MGH Model Shop). The dummy wire was created using tungsten wires (0.00355” outer diameter, Hamilton) with the length matched to the cannula length. The fluorescent dummy wire was created using smaller diameter tungsten wires (0.002” outer diameter, Thermo Scientific Chemicals). 10 mM stock solution of SR101 in DMSO was mixed with a UV curable adhesive (NOA61, Thorlabs). The mixture was applied to the tip of the dummy wire and then cured with an UV LED (M365L3, Thorlabs). The length of the fluorescent dummy wire was matched to the length of the cannula. The schematic for the cannula and design files for the cannula holder and assembly part is available at https://github.com/shou-lab/ShallowAngleCannula.

### Surgery and cannula implantation

Mice were anaesthetized with isoflurane (5% for induction, 1.5% throughout the surgery) using a low-flow anesthesia system (SomnoSuite, Kent Scientific) and placed into a custom head holder (MGH Model Shop). Dexamethasone (0.2 mg kg^−1^) was administered subcutaneously to reduce brain swelling. A heat pad (HTP-1500, Kent Scientific) was used to maintain mouse body temperature during surgery. The scalp was shaved and sterilized. An incision was made to expose the underlying skull. A 1-mm craniotomy was created centered at 2.5 mm lateral to bregma. To ensure, the cannula holder would not make contact with the skull during the implantation process, a rectangular region of skull above the 1-mm craniotomy was progressively thinned down, forming a sloped ramp leading to the 1-mm craniotomy. The mouse’s head position in the holder was adjusted so that the skull at bregma and at 2 mm below bregma were at the same height. The cannula was inserted into its holder and rotated so that the bevel faced towards the top of the brain after implantation. The cannula holder was attached to the manipulator arm of the mouse stereotaxic (RWD Life Science). The angle of manipulator arm was set according to the implantation angle (0). We calculated 0 using the formula, *tan(0) = depth/2mm* where *depth* is the target depth in the brain for the cannula tip.

The mouse was elevated by placing the mouse head holder onto a custom platform. The cannula tip was brought down to touch the mouse dura at the center of the 1-mm craniotomy. The distance (*dist*) that the cannula travels in the brain is calculated using the formula, *dist = sin(0) /depth*. After the cannula is implanted into the brain, the cannula is secured to the skull using dental acrylic (Jet Denture, Lang Dental Manufacturing) and cyanoacrylate glue (Krazy Glue). A cyanoacrylate accelerator (Insta-Set, Bob Smith Industries) is applied to quickly harden the mixture and the holder is retracted. Next, a craniotomy is performed below the location of the implanted cannula by removal of a ∼4 mm piece of the skull. The dura mater was left intact during the craniotomy procedure. A 4-mm-diameter cover glass was placed over the exposed brain and sealed with a mixture of dental acrylic and cyanoacrylate glue. After the cranial window surgery, 0.5 µL of PBS was infused through the cannula and the polyethylene tubing was cut and sealed with dental acrylic and cyanoacrylate glue (see: Cannula infusion section). Mice were given buprenorphine (0.1 mg kg^−1^) and Tylenol for 3 d after surgery, and were allowed to recover for at least 3 weeks before imaging experiments. Prior to imaging, an aluminum headplate with a 5 mm observation hole (CP-1, Narishige Group) was secured to the cranial window using dental acrylic and cyanoacrylate glue.

### Cannula infusion

Mice were anesthetized during the cannula infusion procedure. To prepare for the infusion, polyethylene tubing (PE20, Becton Dickinson) was connected to a 26-gauge hypodermic needle (Becton Dickinson) and a 25 µL syringe (1702 TLLX, Hamilton) was attached to the Luer Lock connector of the needle. The syringe, needle and polyethylene tubing were first filled with PBS and then the tubing was filled with the compound to be infused. A small air gap was introduced between the PBS and compound as a reference to confirm the amount of the compound that entered the brain. To ensure the cannula was clear, the tungsten dummy wire was inserted into and removed from the cannula several times. Then, the open end of the polyethylene tubing was attached to the cannula. A syringe pump (NE-300, New Era Pump Systems) was used to push the syringe for the infusion. The infusion rate for imaging experiments was 0.05 µL/min.

Following infusion of the compound, 0.5 µL of PBS was infused to clear the cannula of the compound. The polyethylene tubing was then cut, and a small drop of dental acrylic and cyanoacrylate glue was applied to the end of the tube to seal it.

### *In vivo* multiphoton microcopy

*In vivo* fluorescence imaging was performed using a commercial multiphoton microscope (FV1000MPE, Olympus). The microscope was equipped with a Ti:sapphire laser (Mai Tai HP DeepSee, Spectra Physics) with tuning range of 690-1040 nm. A 25x water immersion objective (XLPLN25XWMP2, NA = 1.05, Olympus) was used to acquire images. Fluorescence emission was collected in the blue (420-460 nm), green (495-540 nm) and red (575-630 nm) spectral channels. During imaging, mice were anesthetized and body temperature was regulated using a heating pad (Kent Scientific). To stabilize the mouse head for imaging, mice were placed into a headplate holder (MAG-1, Narishige Group). The following volumes and concentrations were used for labeling of various cell types and structures in the brain: 6 µL of 89.0 µM Hoechst 33342 for labeling cell nuclei, 1 µL of 1:5 dilution of stock NeuroTrace 500/525 for labeling pericytes, 1 µL of 20 µM SR101 for labeling astrocytes, 1 µL of 100 µg/ml Isolectin GS-IB4, Alexa Fluor 594 Conjugate for labeling microglia, 1 µL of 100 µM LysoTracker Green DND-26 for labeling lysosomes, 1 µL of 100 µg/ml Fluorescein labeled Wisteria Floribunda Lectin (Vector Laboratories) for labeling PNNs, 2 µL of 1 mg/ml HiLyte Fluor 488-labeled Aβ42 (Anaspec), 1 µL of 100 µg/ml ThioS (Sigma Aldrich) for labeling of amyloid plaques, 2 µL of 5 mg/ml Thiazine Red (Sigma Aldrich) for labeling of NFTs. For angiograms, 100 µL of either 12.5 mg/ml Fluorescein Dextran, 70 kDa or 12.5 mg/ml Rhodamine B Dextran, 70 kDa was injected IV. Fluorescent cell markers were purchased from Thermo Fisher Scientific unless otherwise specified.

### *In vivo* pO_2_ imaging using phosphorescence lifetime imaging microscopy (PLIM)

*In vivo* pO_2_ images were acquired using a custom multiphoton microscope equipped with a tunable pulsed laser (Chameleon Discovery NX, Coherent) (100 fs pulse width, 80 MHz repetition rate). Its output power was modulated by an electro-optic modulator (350-105-02, Conoptics) to generate repeated 300 µs laser excitation cycles (10 µs laser ON time; 290 µs laser OFF time) for pO_2_ imaging or constant average power for angiogram imaging. Lateral imaging was achieved by scanning the laser beam using a two-axis galvanometer mirror system (6210H, Cambridge Technology) with the corresponding scan lens (∼51 mm focal length, 2x AC508-100-B, Thorlabs) and tube lens (180 mm focal length, Olympus). Axial imaging was conducted by moving the objective using a piezo objective scanner (PFM450E, Thorlabs). A 25x water immersion objective (XLPLN25XWMP2, NA = 1.05, Olympus) was used to focus the excitation beam inside the sample. A dichroic mirror (FF875-Di01-38.1×51, Semrock) was used to separate the laser excitation and the emission signals. The emission signals were further filtered by several filters and dichroic mirrors for pO_2_ imaging (FF01-890/SP-50, FF560-Di01-35×52, and FF01-795/150-25, Semrock) and angiogram imaging (FF01-890/SP-50, FF560-Di01-35×52, and FF03-525/50-25, Semrock) before they were detected by two photomultiplier tubes (PMTs) (pO_2_: H10770PA-50, Hamamatsu; Angiogram imaging: R3896, Hamamatsu) respectively. pO_2_ signals were collected using a photon counting unit (C9744, Hamamatsu) and a digital counter (PXIe6341, National Instruments) while the acquisition of angiograms was facilitated by a preamplifier (C9999, Hamamatsu) and an analog input channel (PXIe6356, National Instruments). The whole imaging process was completed under the control of a custom MATLAB software (MathWorks). The temperature of the water immersion was maintained at 36-37ºC using an objective heater (TC-HLS-05, Bioscience Tools) and by applying DC current through insulated nichrome wires (Ace Glass) wrapped around the headplate holder (MAG-A, Narishige Group). The temperature was monitored using a microprobe thermometer (BAT-12, Physitemp Instruments).

To prepare for awake imaging, mice were habituated by being placed in the headplate holder and under the microscope objective for progressively longer durations (10, 20, 30, 45, 60, and 90 minutes) on consecutive days. They received sweetened milk as a reward every 30 minutes during both the training and imaging sessions.

Oxyphor 2P was initially prepared at a stock concentration 80 µM in PBS and was diluted to 20 µM prior to infusion. We infused Oxyphor 2P at 0.05 µL/min, for a total volume of 1 µL, and then infused 0.5 µL of PBS. To generate an angiogram, 100 µL of 12.5 mg/ml Fluorescein Dextran, 70 kDa was injected intravenously. Imaging was performed 2 hours after the infusion. Both Oxyphor 2P and Fluorescein Dextran were excited at 950 nm. Phosphorescence intensity images (only one laser excitation cycle per pixel) were acquired at 250×250 resolution and fluorescent angiograms were acquired at 512×512 resolution. Phosphorescence lifetime data were acquired using 2000 cycles/pixel and in a 20×20 grid. The lifetime of the phosphorescence of Oxyphor 2P was obtained by fitting the phosphorescence decay to a single-exponential function. The lifetime was converted to pO_2_ using a set of parameters determined in independent calibrations (Esipova et al., 2019).

### Immunofluorescence

Mice were euthanized through CO_2_ inhalation and transcardially perfused with 25 mL of PBS followed by 25 mL of 4% PFA using a peristaltic pump (VWR). Brains were extracted and kept in a fixing solution (4% PFA and 15% glycerol in PBS) overnight. Next, brains were dehydrated in sucrose gradients of 10%, 20%, and 30%. After dehydration, brains were sliced into 40 µm coronal sections using a sliding microtome (Leica SM2010R) and sections were stored in 30% glycerol at -20°C until further processing for immunohistochemistry. Sections were subsequently permeabilized with Triton X-100, blocked with normal goat serum (NGS) and incubated for 18 hours with primary antibody IBA1 (rabbit polyclonal anti-IBA1, 1:1000, FUJIFILM Cat# 019-19741, RRID:AB_839504) at 4°C. Sections were washed with PBS and incubated with the secondary antibody (Alexa Fluor 647, 1:500, Molecular Probes Cat# A-21244, RRID:AB_2535812) for 1 h at room temperature. Next, sections were incubated with GFAP (mouse monoclonal anti-GFAP, 1:10000, Sigma-Aldrich Cat# C9205, RRID:AB_476889) for 1 h at room temperature. After washing, sections were cover slipped with antifade mounting medium with DAPI (Vector Laboratories, H-1500) and imaged by confocal microscopy (Olympus FV3000).

## ACKNOWLEDGMENTS

We thank Buyin Fu for technical support. This study was supported by NIA K01AG072046, NIBIB U24EB028941, NIA R01AG054598 and a Heitman Young Investigator Career Development Award. The content expressed is solely the responsibility of the authors and do not necessarily represent the official views of the NIH.

## CODE AND DATA AVAILABILITY

The data that support the findings of this study are available from the authors upon reasonable request. See author contributions for specific data sets.

## AUTHOR CONTRIBUTIONS

S.S.H., J.Y., and B.J.B. conceived the study and designed the experiments. S.S.H. and Y.T. performed the immunostaining and confocal imaging of tissue sections, S.S.H., J.Y., Y.K., C.D. and M.C.R. performed the *in vivo* validation experiments, and analyzed the results, S.S.H., Y.K. and C.D. performed the tracking of degenerating neurons in AD mice, M.E.K. and S.A.V. synthesized the pO_2_ probe and performed analysis and interpretation of pO_2_ data, S.S.H., Y.K., Q.P., and S.S. performed the *in vivo* longitudinal pO_2_ imaging and interpreted the results. S.S.H. and B.J.B wrote the manuscript. All authors have given approval to the final version of the manuscript.

