## Supplementary figures for "Shallow-angle intracranial cannula for repeated infusion and in vivo imaging with multiphoton microscopy"

**SUPPLEMENTARY INFORMATION**

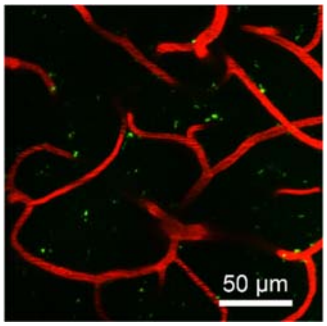


**Figure S1 | Fluoro Jade C infusion in rTg4510 wild-type littermates.** FJC (green) was administered through the cannula system while Rhodamine B Dextran (red) was injected IV to generate an angiogram. The representative image from wild-type littermate mice (n = 3) shows absence of FJC- positive cells in the imaging region adjacent to the cannula. No FJC-positive cells were detected in any of the imaged wild-type mice.

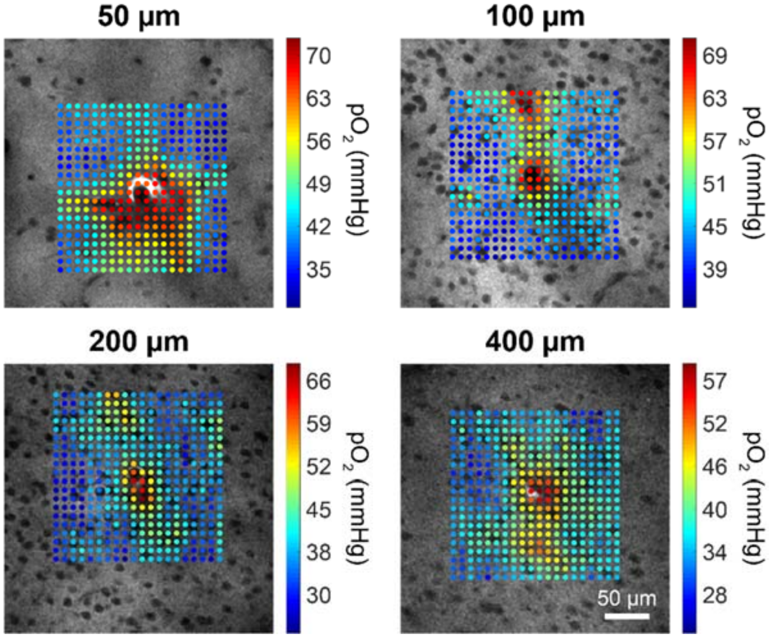


**Figure S2 | *In vivo* imaging of brain tissue $pO_2$ in anesthetized mice.** C57BL/6J mice (n = 2) were infused with Oxyphor 2p and imaged under 1% isoflurane. Representative images showing color-coded $pO_2$ values overlaid onto phosphorescence intensity images. Images were acquired at 50, 100, 200 and 400 µm from the brain surface.

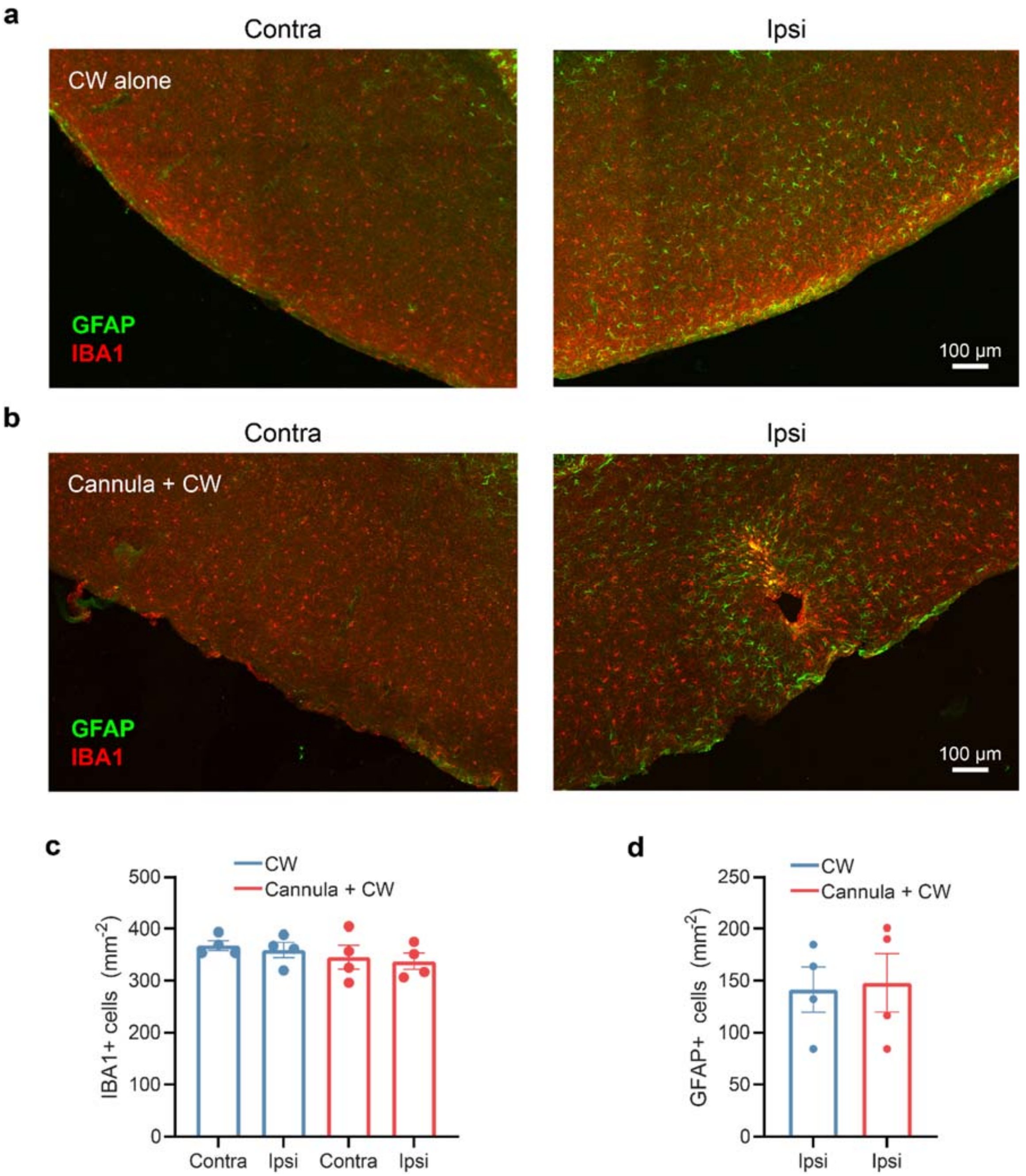


**Figure S3 | Immunohistochemical characterization of the neuroinflammatory response due to chronic cannula implantation.** Cranial window (CW) surgery (n = 4) or cannula implantation followed by CW surgery (n = 4) were performed on C57BL/6J mice. Mice were sacrificed 30 days post-surgery, and brain sections were immunostained for microglia (anti-IBA1) and activated astrocytes (anti-GFAP). Representative images show contralateral and ipsilateral hemispheres from mice with (**a**) CW and (**b**) cannula + CW surgeries. Quantitative analysis of the number of (**c**) IBA1 and (**d**) GFAP positive cells is shown for both CW and cannula + CW mice.
